# TopFD - A Proteoform Feature Detection Tool for Top-Down Proteomics

**DOI:** 10.1101/2022.10.11.511828

**Authors:** Abdul Rehman Basharat, Yong Zang, Liangliang Sun, Xiaowen Liu

## Abstract

Top-down liquid chromatography-mass spectrometry (LC-MS) analyzes intact proteoforms and generates mass spectra containing peaks of proteoforms with various isotopic compositions, charge states, and retention times. An essential step in top-down MS data analysis is proteoform feature detection, which aims to group these peaks into peak sets (features), each containing all peaks of a proteoform. Accurate protein feature detection enhances the accuracy in MS-based proteoform identification and quantification. Here we present TopFD, a software tool for top-down MS feature detection that integrates algorithms for proteoform feature detection, feature boundary refinement, and machine learning models for proteoform feature evaluation. We performed extensive benchmarking of TopFD, Promex, FlashDeconv, and Xtract using five top-down MS data sets and demonstrated that TopFD outperforms other tools in feature accuracy, reproducibility, and feature abundance reproducibility.

## 1. Introduction

Top-down mass spectrometry (MS) has attracted increasing attention owing to its unique capacity to analyze intact proteoforms and characterize proteoforms with multiple alterations [1-3]. Advances in high resolution and high accuracy MS instruments significantly increased proteoform identifications and amino acid sequence converge in proteome-wide top-down proteomics analysis [4]. Recent top-down MS studies identified more than 23,000 proteoforms from colorectal cancer cells [5] and about 30,000 proteoforms from human blood and bone marrow cells [6]. Top-down MS-based proteoform profiling has successfully identified differentially expressed proteoforms associated with diseases [7-10].

Proteoform feature detection is a fundamental computational problem in top-down MS-based proteoform quantification. In proteome-wide top-down MS analysis, proteoforms extracted from samples are first separated by liquid chromatography (LC) or other separation methods and then analyzed by tandem mass spectrometry (MS/MS). Each proteoform has an elution profile (Fig. 1a), which depicts the abundance of the proteoform eluted over time in proteoform separation. A mass spectrum contains a list of peaks, each represented by its mass-to-charge ratio (*m*/*z*) and intensity. The isotopologues of a proteoform with the same charge state are detected as a group of isotopic peaks in a mass spectrum, called an isotopic envelope (Fig. 1b). The peak intensities in an isotopic envelope follow a distribution determined by the isotopic frequencies of the atoms in the proteoform. A mass spectrum often contains multiple isotopic envelopes of a proteoform with different charge states (Fig. 1b). Feature detection in top-down MS aims to identify all isotopic peaks of each proteoform over retention time (RT) and across charge states in an LC-MS data file and reports its elution profile and total signal intensity (Fig. 1c).

**Figure 1.**
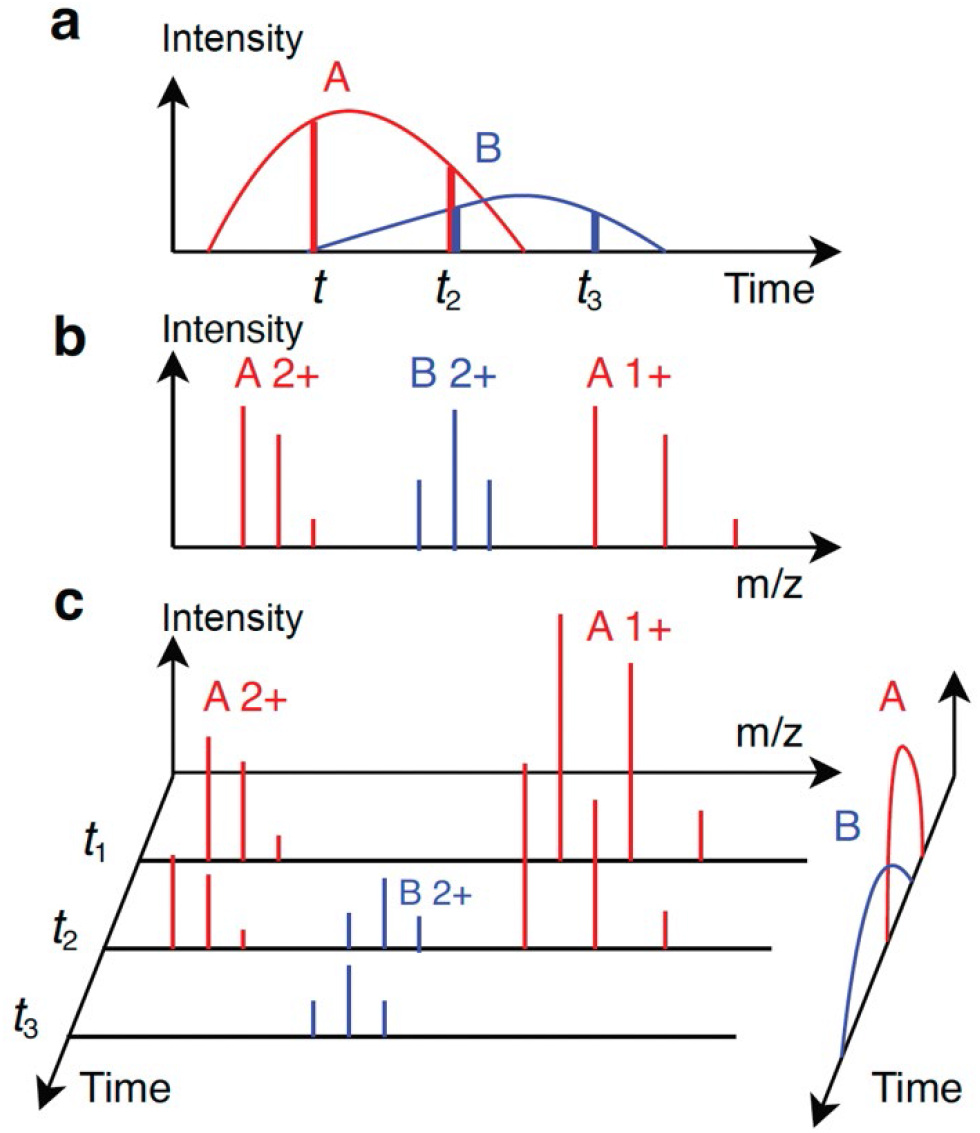
An illustration of two proteoform features in an LC-MS map. (a) Elution profiles (curves) of two proteoforms *A* and *B*. At time points *t*_1_, *t*_2_, and *t*_3_, three MS1 spectra are generated. The abundances of *A* have a ratio of 4:3:0 (two red vertical lines), and the abundances of *B* have a ratio of 0:1:1 (two blue vertical lines). (b) Theoretical isotopic envelopes of proteoform *A* with charge states 1+ and 2+ and proteoform *B* with charge state 2+. (c) Three MS1 spectra are generated at time points *t*_1_, *t*_2_, and *t*_3_. The elution profiles on the right are the same as subfigure (a). The three spectra contain four experimental isotopic envelopes of *A* and two envelopes of *B*.

Many methods have been proposed [11-22] for peptide feature detection in bottom-up MS (Supplementary Table S1), which is similar to proteoform feature detection in top-down MS. These methods are focused on solving three computational problems in feature detection: (1) grouping peak signals with similar *m*/*z* values in consecutive MS1 scans into an *m*/*z* or envelope trace (Fig. 1c), (2) splitting a trace into several ones corresponding to single peptides if the trace contains peak signals from two or more peptides, and (3) evaluating and ranking reported features. To identify an *m*/*z* or envelope trace, a seed peak or envelope of a feature and its corresponding scan are first selected, and then the peak or envelope is extended along the RT in both directions until one or several scans do not contain peaks or envelopes with similar *m*/*z* values matched to the seed [12].

Trace splitting methods can be divided into three groups. The first is to split a trace at the scans lacking matched peaks or envelopes [12]. In the second approach, the elution profile of a feature is fitted to a distribution or function, such as a Gaussian distribution or wavelet function, and a cut-off signal intensity is used to determine feature boundaries [15, 23]. The third approach is to use a function, such as a Savitzky-Golay filter [16, 19, 22], to smooth a trace locally and use local minima to determine feature boundaries in the trace.

Reported features are in general evaluated by their peak intensities, peak *m*/*z* errors, and RT ranges [21]. The quality of envelope features is also determined by the similarity of theoretical and experimental isotopic peak intensity distributions [12]. Recently, deep learning methods have been proposed to evaluate envelope features [24].

Feature detection in top-down MS is more challenging than in bottom-up MS, as top-down mass spectra tend to have higher charge state ions, more complex isotopic envelopes, and more overlapping envelopes than bottom-up spectra. As a result, feature detection methods designed for bottom-up MS may fail to achieve good performance for top-down MS.

Several methods have been proposed for feature detection in top-down MS, e.g., Xtract [25, 26], ProMex [27], and FlashDeconv [28]. ProMex uses a greedy algorithm to cluster isotopic envelopes of the same proteoform across MS1 scans. The peak intensities in the experimental envelopes of a proteoform are aggregated to reduce the measurement errors between theoretical and experimental isotopic distributions. The elution profile of each proteoform feature is constructed and smoothed by a Savitzky-Golay filter. Finally, the RT range is obtained using 1% of the apex intensity as the signal intensity cutoff. The quality of each feature is evaluated by a likelihood ratio function based on a Bayesian network model. In FlashDeconv [28], candidate features are identified by searching a mass spectrum for peak groups that are generated from proteoform molecules with the same mass and different charge states. The RT range of a feature is determined by a mass trace detection algorithm, in which features are extended along RT and feature boundaries are found using a smoothing method [29]. A feature is evaluated by fitting a Gaussian distribution to the peak intensities with different charge states and computing the cosine similarity between the fitted and experimental intensities. The methods in Xtract have not been published.

In this paper, we propose TopFD, a method for proteoform feature detection in top-down MS, in which the functions in MS-Deconv [30] are employed to identify feature candidates, and RT boundaries of feature signals are determined using local minima of envelope traces. In addition, a neural network model that takes eight attributes of proteoform features as the input was trained for feature evaluation. TopFD was extensively assessed and compared with ProMex [27], FlashDeconv [28], and Xtract using five top-down MS data sets. Experimental results demonstrated that TopFD outperforms these tools in the accuracy and reproducibility of proteoform feature detection and the reproducibility of proteoform quantification.

## 2. Results

### 2.1 Proteoform feature detection and evaluation

In an LC-MS experiment, the start, apex, and end RTs of a proteoform are determined by the separation column, the experimental parameters, and the chemical and physical properties of the proteoform. An MS1 spectrum collected during the RT range of a proteoform contains peaks of the proteoform, which can be grouped into one or several isotopic envelopes based on their charge states (Fig. 1b). The set of all isotopic envelopes of a proteoform with a specific charge state in the LC-MS map is an *envelope set* (single charge feature) of the proteoform. The collection of the envelope sets of the proteoform for all charge states is the *envelope collection* (multi-charge feature) of the proteoform. The proteoform feature detection problem aims to find all envelope collections in an LC-MS map and report the monoisotopic mass, RT range, and abundance of each envelope collection (Fig. 1c).

Fig. 2 shows the overall scheme of TopFD for proteoform feature detection (Methods). In preprocessing, TopFD filters out noise peaks in an LC-MS map and then uses the functions in MS-Deconv [30] to identify experimental isotopic envelopes of proteoforms in single MS1 spectra. A theoretical isotopic envelope is computed for each experimental isotopic envelope using the Averagine model [31]. The reported isotopic envelopes are ranked based on their total peak intensities and then used iteratively as seed envelopes for feature detection.

**Figure 2.**
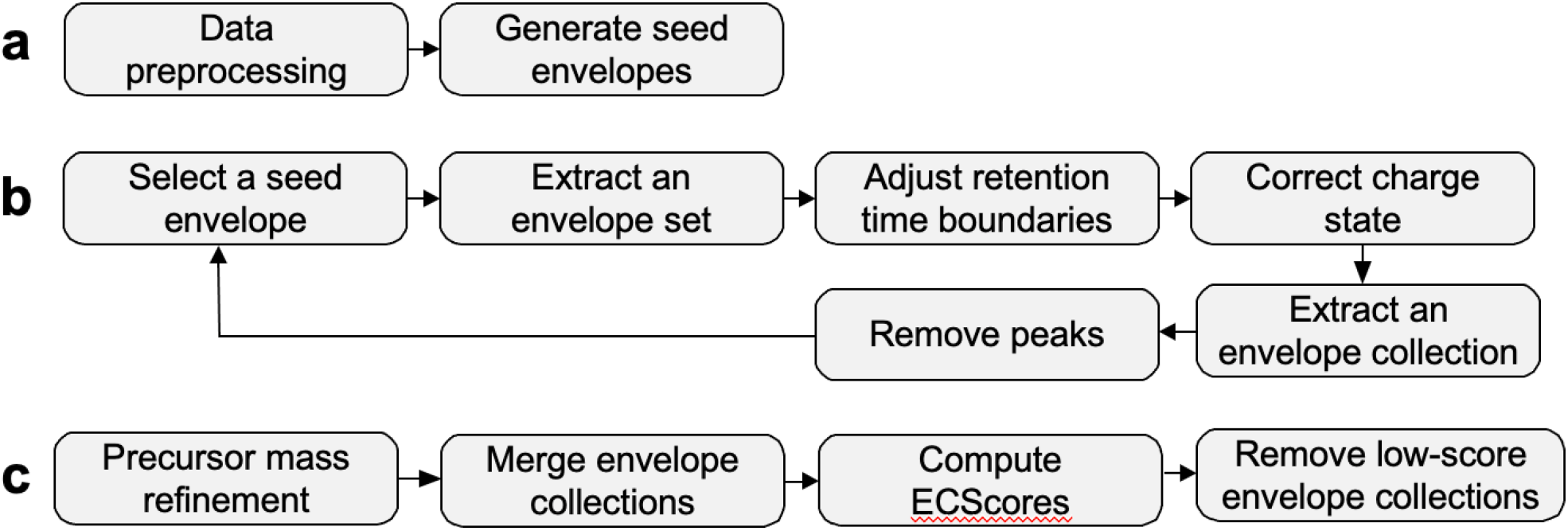
An overview of the pipeline for proteoform feature identification in TopFD. (a) Preprocessing. Experimental centroided peaks are processed to remove those that have a low intensity or appear in only one MS1 spectrum. Then MS-Deconv is used to deconvolute MS1 spectra to obtain seed envelopes. (b) Proteoform feature exaction. (1) The reported seed envelopes are ranked based on the sum of the peak intensities of the theoretical envelope. The one with the highest intensity is selected. (2) To extract an envelope set, peaks in the seed theoretical envelope are matched with experimental peaks and extended in both forward and backward directions until no matching experimental peaks are found. (3) The RT boundaries of the reported envelope set are refined if it contains peaks from neighboring envelope sets. (4) The charge state of the envelope set is evaluated and corrected if needed. (5) Once an envelope set is extracted, the neighboring charge states are explored to find other envelope sets in the envelope collection. (6) The experimental peaks included in the envelope collection are removed from the data. The six steps are repeated for the next seed envelope, which has the highest intensity in the remaining seed list. (c) Postprocessing. The precursor masses of reported envelope collections are first refined. Envelope collections are then merged if they have similar precursor masses and similar retention time ranges. Finally, an ECScore is computed for each envelope collection and those with low ECScores are removed.

In feature detection, TopFD first extends a seed envelope to neighboring MS1 scans to obtain an envelope set, then the RT boundaries and charge state of the envelope set are adjusted (Methods). Next, the envelope set is extended to identify envelope sets with the same precursor mass and neighboring charge states, resulting in an envelope collection. Finally, peaks used in the envelope collection in the LC-MS map will be removed or reduced.

In postprocessing, the precursor mass of each envelope collection is refined using its isotopic peaks, and then envelope collections with similar precursor masses and RTs are merged. Finally, a neural network model with four fully connected layers (Methods) is used to assign an envelope collection score (ECScore) to each envelope collection, and those with low ECScores are removed.

### 2.2 Training the neural network model for ECScore

A set of envelope collections for training ECScore was generated from three top-down data sets: one from SW480 cells with technical triplicates and the other two from breast cancer (BC) samples, each with six replicates (Methods). On average, 21,190 envelope collections were reported from each SW480 replicate and 1,765 from each BC replicate (Supplementary Table S2) using the methods for envelope collection identification and merging (Fig. 2) with the default parameter settings (Supplementary Table S3). Note that all envelope collections reported from the three data sets were used for generating training data. We labeled the envelope collections identified from the first SW480 replicate, and the first replicate of each BC data set as follows: An envelope collection was labeled negative if it was reported in only the first replicate and labeled positive if it was reported in all the three SW480 replicates or ≥ 5 BC replicates. The unlabeled envelope collections were removed. A total of 8,579 envelope collections were labeled positive, and 10,876 were negative. We randomly split the envelope collections with a 67:33 ratio into training and validation sets and trained the neural network model for ECScore (Methods). ECScore achieved a balanced accuracy of 87.03% and the area under the receiver operating characteristic (ROC) curve (AUC) value of 94.18% on the validation data set. The default cutoff of ECScore was set to 0.5 for filtering out low-quality envelope collections because the ECScore distributions of the validation envelope collections show that the cutoff value of 0.5 can separate positive envelope collections from negative ones (Supplementary Fig. 1).

### 2.3 Comparison of ECScore and EnvCNN

We compared the accuracy of ECScore and the EnvCNN score [32] on two top-down MS data sets: one from SW620 cells with three replicates and the other from ovarian cancer (OC) samples with ten replicates (Methods). Similar to the training data, we labeled the envelope collections reported from the first replicates of the SW620 and OC data. An envelope collection was labeled negative if it was reported in only one replicate and labeled positive if it was reported in all three SW620 replicates or ≥ 8 OC replicates. This resulted in an SW620 test set of 7,446 positive and 2,862 negative envelope collections and an OC test set of 5,682 positive and 286 negative envelope collections. Because the EnvCNN model takes single isotopic envelopes, not envelope collections, as the input, all test envelope collections were converted to aggregate experiment envelopes (Section 4.6 in Methods), which were used as the input of the EnvCNN model.

ECScore achieved higher ROC AUC values (Supplementary Fig. 2) than the EnvCNN score on the OC test set (92.79% vs 81.08%) and the SW620 test set (79.36% vs 61.48%). We also compared the rank-sum values of the two scoring functions on the OC and SW620 test sets. To compute the rank-sum of a list of envelope collections, all envelope collections were ranked in the decreasing order of their scores, and the ranks of all positive envelope collections were summed up. ECScore reduced the rank-sum values compared with the EnvCNN score on the OC data set (1.68 ×10^8^ vs. 1.85 ×10^8^) and the SW620 data set (1.11 ×10^8^ vs. 1.30 ×10^8^).

### 2.4 Evaluation of the artifacts of reported proteoform features

Following the methods in Jeong *et al*. [28], we assessed the quality of proteoform features reported by feature detection tools using three types of artifact masses: low harmonic masses, high harmonic masses, and isotopologues. Incorrect charge state assignments to isotopic envelopes will result in low and high harmonics masses, which are integer fractions and multiples of true masses of proteoforms, respectively. Errors in computing the monoisotopic masses of envelope collections will introduce isotopologues, which are shifted by the mass of one or several neutrons compared with true masses.

An envelope collection *A* is a mass artifact of another envelope collection *B* if (1) the total peak intensity of *B* is higher than *A*, (2) the overlapping RT range of *A* and *B* is larger than 80% of the RT range of *A*, and (3) the monoisotopic mass of *A* is an isotopologue, low harmonic mass, or high harmonic mass of the monoisotopic mass of *B* (Methods). An envelope collection is *valid* if it is not a mass artifact of another envelope.

We benchmarked TopFD, ProMex (version 1.1.8082) [27], FlashDeconv (version 2.0) [28], and Xtract (Thermo BioPharma Finder 4.1) [25, 26] in the ratio of valid proteoform features using the first OC replicate and the first SW620 replicate. Parameter settings, running times, and numbers of reported proteoform features of the tools are given in Supplementary Tables S3-S6, Fig. S3, and Supplementary Table S7, respectively. For each tool, we ranked the reported proteoform features based on their total peak intensities and plotted the ratio of the valid ones against the number of top features. We chose total peak intensities, not software tool-specific scores, to rank features to ensure a fair comparison of the tools. Proteoform features reported by TopFD achieved the best valid ratios (>82%) among the four tools (Figs. 3a, 3d). The valid ratios for Xtract and FlashDeconv are also high, and distributions of the three types of artifacts are similar for TopFD, FlashDeconv, and Xtract (Figs. 3b, 3c, 3e, 3f). ProMex reported many isotopologues, resulting in low valid percentages.

**Figure 3.**
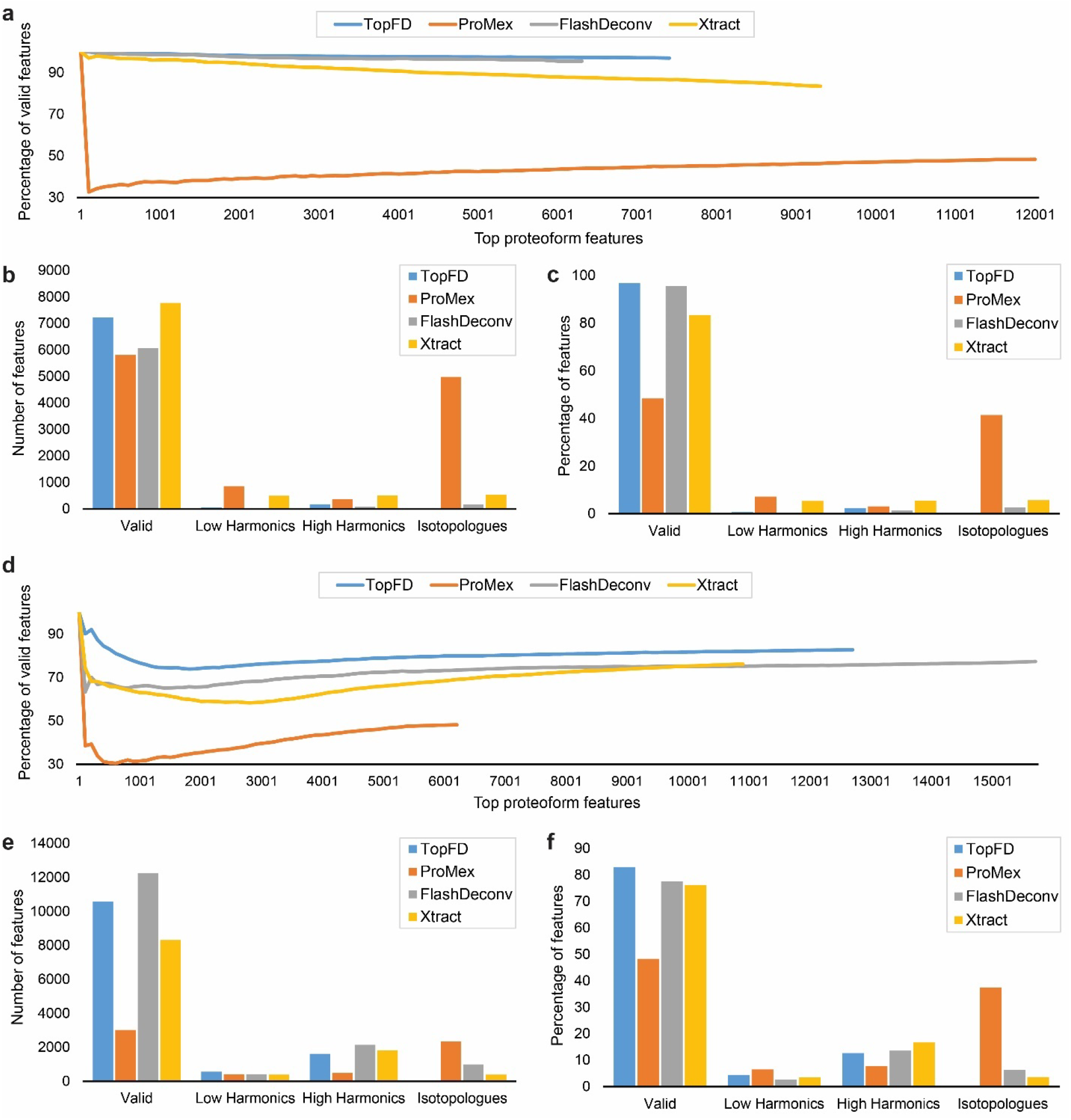
Artifact masses reported by TopFD, ProMex, FlashDeconv, and Xtract from the first replicate of the OC data and the first replicate of the SW620 data. (a) The number of top proteoform features against the percentage of valid features for the first replicate of the OC data. Numbers (b) and percentages (c) of valid, high harmonic, low harmonic masses, and isotopologues in all features reported by the tools from the first replicate of the OC data. (d) The number of top proteoform features against the percentage of valid features for the first replicate of the SW620 data. Numbers (b) and percentages (e) of valid, high harmonic, low harmonic masses, and isotopologues in all features reported by the tools from the first replicate of the SW620 data.

### 2.5 Comparison between total ion currents (TICs) and feature intensities

While the TIC of an MS1 scan depicts the number of ions detected by the scan, the total peak intensity of reported proteoform features for an MS1 scan gives the number of ions reported by feature detection tools. These two measurements are expected to be consistent with each other. So following the method in [28], we compared the TICs and total feature intensities reported by the four tools.

To evaluate the correlation between TICs and proteoform feature intensities, we divided the RT range of an MS data set into 1-minute RT bins and computed the TIC and total proteoform feature intensity for each bin. The TIC of an RT bin is the sum of the total TIC of all MS1 scans in the bin; the total proteoform feature intensity of an RT bin is the sum of the intensities of the proteoform features whose apex RTs are in the bin. All mass artifacts were removed before the computation of total proteoform feature intensities. The total feature intensities reported by TopFD achieved the best similarity with the TICs on the OC and SW620 data sets (Fig. 4). For the first OC replicate, the cosine similarity scores between the two measurements are 78.04%, 77.46%, 77.40%, and 61.37% for TopFD, ProMex, FlashDeconv, and Xtract, respectively. Similarly, for the first SW620 replicate, the cosine similarities are 74.65%, 23.61%, 73.68%, and 67.67% for TopFD, ProMex, FlashDeconv, and Xtract, respectively (Supplementary Fig. S4).

**Figure 4.**
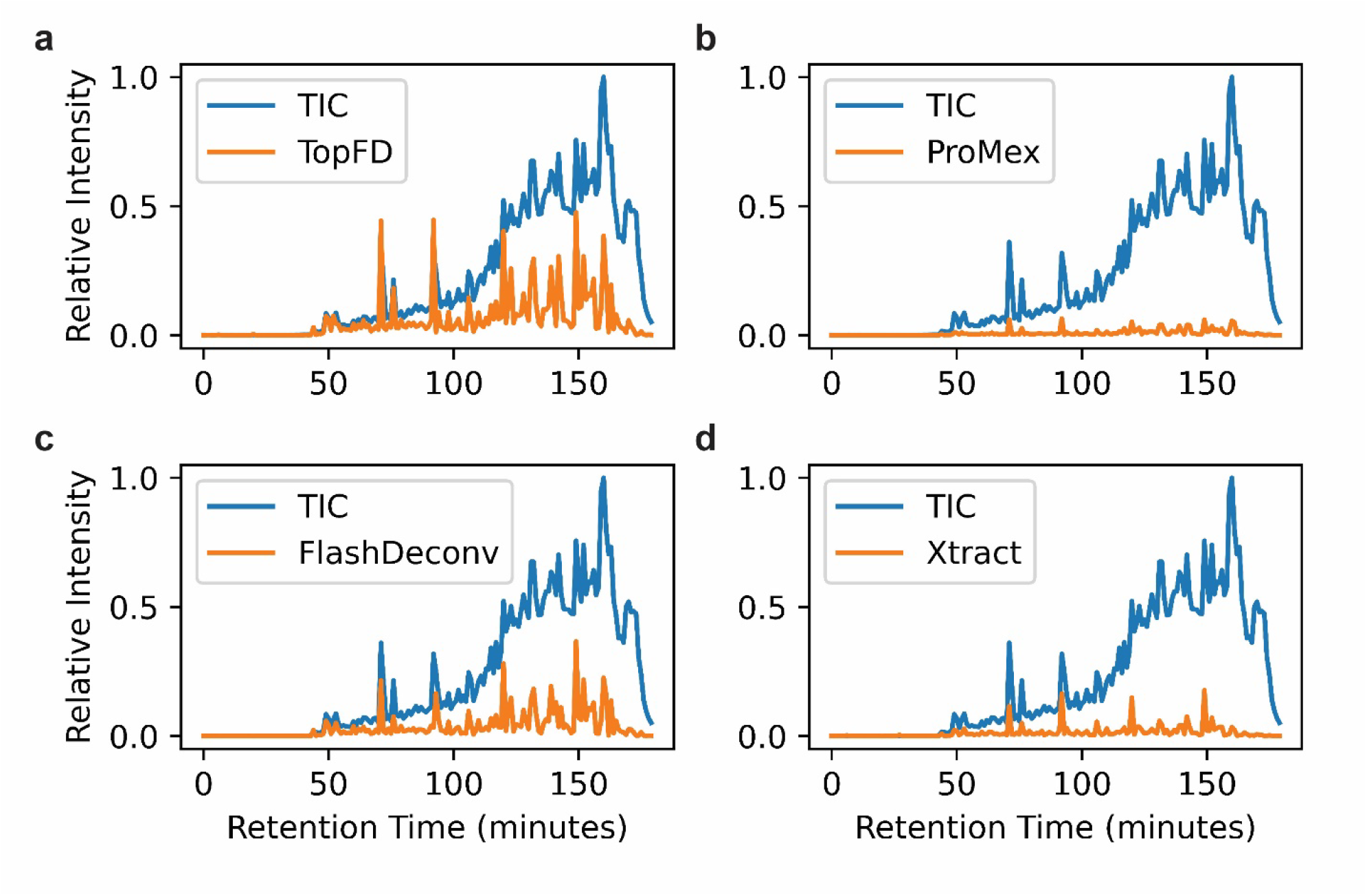
Comparison of TIC and total proteoform feature intensities reported by feature detection tools along the RT time for the first OC replicate. (a) TopFD, (b) ProMex, (c) FlashDeconv, and (d) Xtract. The RT range of the MS data is divided into 1-minute RT bins. The TIC and total proteoform feature intensity in each bin are normalized by dividing them by the maximum TIC value.

### 2.6 Reproducibility of proteoform features in MS replicates

We benchmarked the feature reproducibility of TopFD against ProMex, FlashDeconv, and Xtract on the OC and SW620 data sets. In MS technical replicates, a true proteoform feature is expected to be observed in all the replicates, so the frequencies of proteoform features reported from MS technical replicates are a good metric for evaluating proteoform features [27]. Because mass artifact removal can improve the quality of reported proteoform features, we removed mass artifacts from proteoform features reported by the tools. The four tools reported different numbers of proteoform features from an MS data file, so we kept the top *n* features in each of the four feature lists to ensure a fair comparison, where *n* is the minimum size of the four feature lists. On average, 6,034 and 3,048 features were kept for each replicate of the OC and SW620 data sets, respectively. For each tool, the features reported from the first replicate (5,811 for OC and 3,025 for SW620) were compared with those reported from other replicates to obtain their numbers of occurrences.

TopFD reported the highest number (3,410 out of 5,811) of proteoform features reported in all ten replicates of the OC data set (Fig. 5a) and the highest percentage (76.80%) of proteoform features in 8 or more replicates compared with ProMex (65.96%), FlashDeconv (57.76%), and Xtract (62.14%). Similarly, TopFD outperformed the other tools in feature reproductivity on the SW620 data set (Fig. 5b). A total of 65.32% proteoform features reported by TopFD were observed in all three replicates, which was better than ProMex (57.45%), FlashDeconv (47.80%), and Xtract (55.63%). The performance of TopFD on the two data sets demonstrates its high reproducibility, which also provides evidence for its high accuracy.

**Figure 5.**
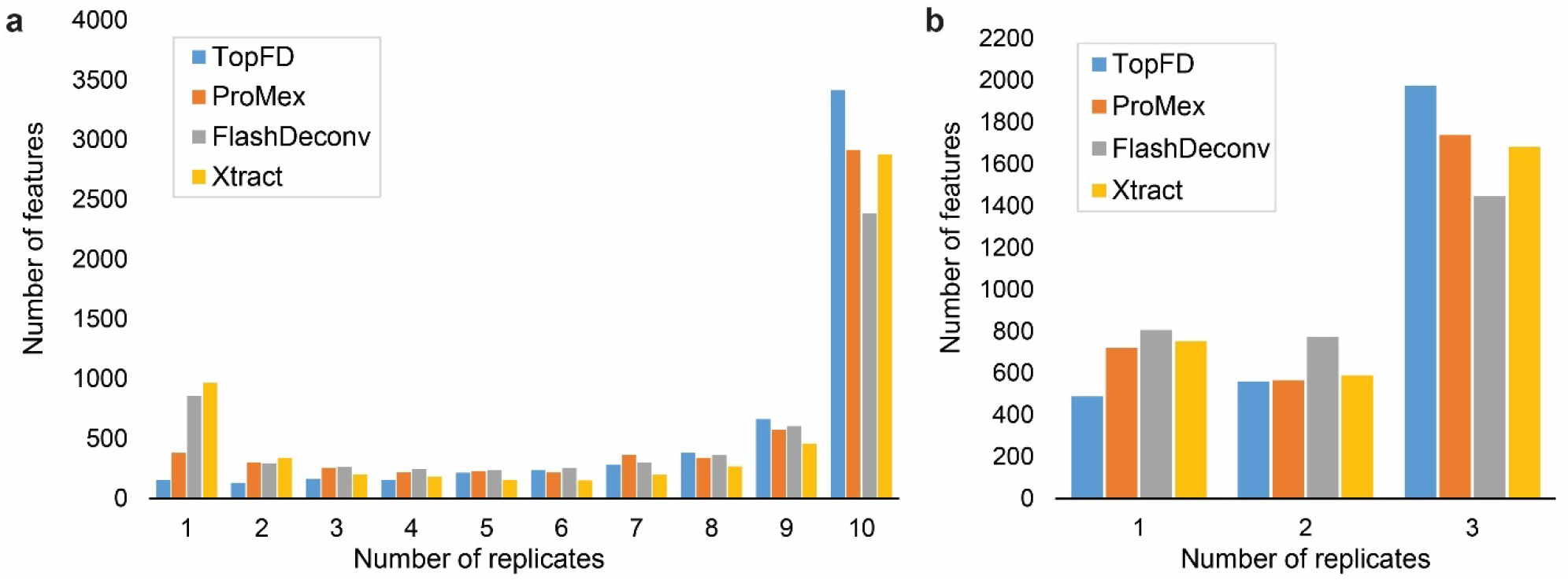
Comparison of the reproducibility of proteoform features reported by TopFD, Promex, FlashDeconv, and Xtract in MS replicates. (a) The frequencies of feature observations in the OC data set for the 5,881 features reported from the first replicate. (b) The frequencies of feature observations in the SW620 data set for the 3,048 features reported from the first replicate.

### 2.7 Quantitative Reproducibility

High-accuracy proteoform feature detection is essential for increasing the reproducibility of proteoform abundances measured in MS replicates. We benchmarked the four tools in the reproducibility of proteoform abundances using the OC and SW620 data sets. Mass artifacts were removed, and only proteoform features identified in all the replicates were kept. The log abundances of the proteoforms were obtained for each replicate, and the Pearson correlation coefficient (PCC) was computed for each pair of replicates. TopFD reported better proteoform abundance reproducibility compared with other tools, thus indicating high reproducibility in proteoform quantification on the OC (Fig. 6) and SW620 (Supplementary Fig. S5) data sets.

**Figure 6.**
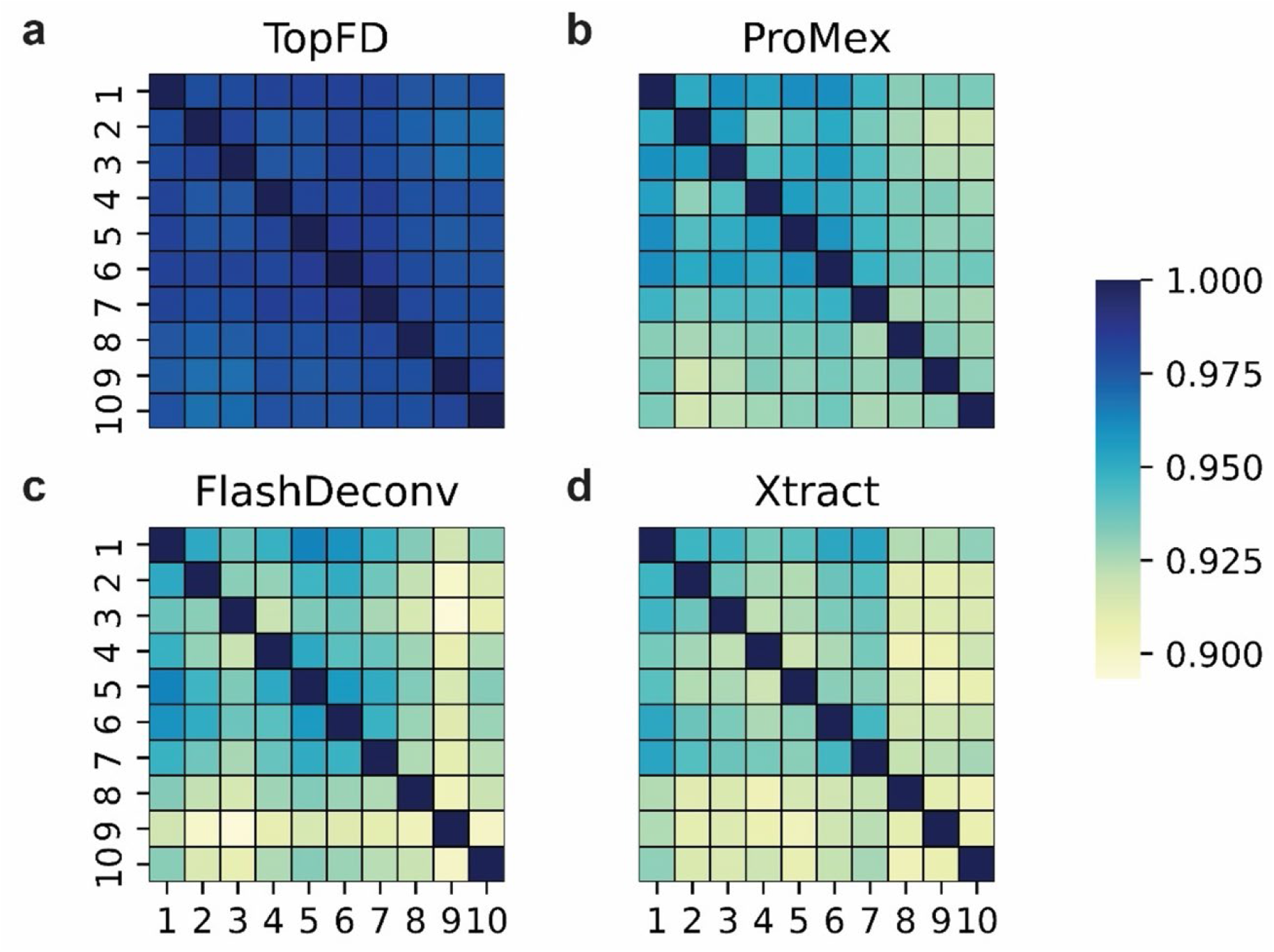
Quantitative reproducibility of proteoform features reported from the OT data set. The PCC between the abundances of proteoform features is obtained for each replicate pair for a) TopFD, b) ProMex, c) FlashDeconv, and d) Xtract.

## 3. Discussion

Accurate proteoform feature detection in top-down MS is essential for proteoform quantification. Proteoform feature detection results show that more than 58% of features reported from one replicate are observed in all three replicates of the SW620 data and all ten replicates of the OT data, and the PCCs of proteoform abundances between MS replicates are higher than 93%. This level of reproducibility makes it possible to identify differentially expressed proteoforms in two types of samples in proteome-wide top-down MS studies.

Accurate precursor monoisotopic mass calculation in top-down MS is indispensable for identifying PTMs and other alterations in proteoforms. Proteoform feature detection tools often report monoisotopic masses of features with +/-1 Da errors due to noise in measured isotopic peak intensities in experimental envelopes and errors in isotopic peak intensities in theoretical envelopes. Following the method in Park et al. [27], TopFD aggregates isotopic envelopes in an envelope collection to improve the accuracy of experimental isotopic peak intensities. The errors in theoretical peak intensities are mainly introduced by the difference between the chemical composition of the proteoform and that computed based on the Averagine model [31]. To address this problem, a post-processing step can be employed to recalculate the theoretical isotopic peak intensities when the proteoform sequence of the feature is identified and its chemical composition is known.

ECScore in TopFD outperforms the EnvCNN score for computing confidence scores for identified envelope collections, showing that neural network-based models have the potential to improve the accuracy in proteoform feature detection. ECScore is based on a simple fully connected neural network with eight attributes of envelope collections as the input. As complex neural network models [24] have been successfully used for peptide feature detections in bottom-up MS, a future research direction is to employ deep learning models to solve various problems in proteoform feature detection, such as feature boundary detection and scores of envelope collections, and use these models to further improve the accuracy in proteoform feature detection. There are still many challenging problems in proteoform feature detection, like feature boundary detection and the identification of overlapping proteoform features and low abundance features. TopFD identifies overlapping proteoform features by comparing isotopic peak intensities in experimental and theoretical envelopes. If there is a significant difference between the intensities of a pair of matched experimental and theoretical peaks, the experimental peak is treated as an overlapping peak. TopFD relies on seed envelopes for proteoform feature identification and may fail to find seed envelopes for low abundance features. Deep learning models are promising to provide better solutions for identifying overlapping features and low abundance features.

## 4. Methods

### 4.1 Data Sets

Five top-down MS data sets were used in this study, and msConvert [33] was used to convert raw files into centroided mzML files. The first two data sets were generated from SW480 and SW620 colorectal cancer (CRC) cells and are available at MassIVE (ID: MSV000090488). Proteoforms extracted from the sample were analyzed using a Thermo Q-Exactive HF mass spectrometer coupled with a 105-minute nanoRPLC separation system. The top 5 precursor ions in each MS1 spectrum were selected from an isolation window of 4 *m*/*z* for MS/MS analysis using higher-energy collisional dissociation (HCD). Both MS1 and MS/MS spectra were acquired at a resolution of 120,000 (at 200 *m*/*z*). Three technical replicates were obtained for each cell line.

The third data set was generated from ovarian cancer (OC) samples [27] and downloaded from MassIVE (ID: MSV000080257). In the experiment, five OC patient samples were pooled, and the extracted proteoforms were analyzed using a Thermo Velos Orbitrap Elite mass spectrometer coupled with a 180-min LC separation system. The top 4 precursor ions in each MS1 spectrum were selected separately with an isolation window of 10 *m*/*z* for MS analysis using collision-induced dissociation (CID). MS1 and MS/MS spectra were acquired at a resolution of 240,000 (at 400 *m*/*z*) and 120,000 (at 400 *m*/*z*), respectively. A total of 10 MS experiment replicates were obtained.

The fourth and fifth data sets were generated from two patient-derived mouse xenografts derived from basal-like and luminal-B human breast cancer samples [34], which were downloaded from the CPTAC data portal [35]. The GELFrEE method [36] was performed for each sample to obtain a fraction containing proteoforms of size up to 30 kDa. Subsequently, six technical replicates were generated for each sample. The samples were analyzed using a Thermo Orbitrap Elite mass spectrometer coupled with an LC system with 90-minute separation. The top two precursor ions in each MS1 spectrum were selected from an isolation window of 15 *m*/*z* for MS/MS analysis using higher-energy collisional dissociation (HCD). MS1 and MS/MS spectra were acquired at a resolution of 120,000 (at 400 *m*/*z*) and 60,000 (at 400 *m*/*z*), respectively.

### 4.2 Peak filtering

Two methods were used to filter out noise peaks in MS1 spectra to speed up proteoform feature detection. The first filtering method was based on peak intensities, in which peaks with intensity lower than a cutoff intensity were removed because most of them do not provide valuable information for feature detection. To obtain the cutoff intensity, a histogram of the intensities of all MS1 peaks in the data file was generated, and the noise intensity level, denoted by *h*, was set to the middle value of the bin with the highest frequency [30], and the cutoff intensity was set to 3*h*. The second filtering method was based on the number of consecutive spectra in which a peak is observed. Peaks that appear in only one MS1 spectrum, not several consecutive MS1 spectra, tend to be noise ones. Therefore, a peak in an MS1 spectrum was removed if it was not observed in its neighboring MS1 scans within an *m*/*z* error tolerance of 0.01.

### 4.3 Seed envelope identification

We obtained isotopic envelopes of proteoforms from single spectra and then used them as seeds to find envelope sets and envelope collections. Experimental isotopic envelopes in single MS1 spectra were identified based on the methods in MS-Deconv [30, 37] with eight steps. (1) A peak in the spectrum is selected as the base peak of the envelope. (2) A theoretical isotopic distribution is computed using the Averagine model [31] with a given charge state so that the *m*/*z* value of the highest intensity peak in the envelope equals the *m*/*z* value of the base peak. (3) Peaks in the theoretical distribution are matched to those in the spectrum by comparing their *m*/*z* values with an error tolerance (0.02 in the experiments). The set of matched experimental peaks is reported as an experimental isotopic envelope. (4) A theoretical envelope is obtained by scaling the peak intensities of theoretical distribution so that the sum of the intensities of the top three peaks in the theoretical envelope is the same as that of the top three experimental peaks. (5) The theoretical and experimental envelope pair is scored using the default scoring function in MS-Deconv, and its monoisotopic mass is computed. (6) Peaks in the envelope pair are removed if their scaled theoretical intensities are lower than a cutoff intensity, which is set to the intensity of *3h* (Section 4.2). (7) After all candidate envelopes are generated from the spectrum, a dynamic programming method is used to report a group of theoretical and experimental envelope pairs that fits the spectrum. (8) The envelope pairs are further filtered using a cutoff value (0.5 in the experiments) for the PCC between the peak intensities of theoretical and experimental envelopes.

### 4.4 Extending seed envelopes to envelope sets

We ranked the experimental and theoretical envelope pairs reported from all MS1 spectra in an LC-MS run in the decreasing order of the total peak intensity, which is the sum of the peak intensities of the theoretical envelope. The theoretical envelope with the highest intensity was selected as the first seed envelope, which was then extended to neighboring scans to obtain an envelope set. Theoretical envelopes were used for the extension because they tend to have fewer errors in *m*/*z* values and peak intensities than experimental envelopes.

To obtain an envelope set, a seed envelope *E* of a proteoform was matched to experimental peaks in its neighboring spectra to extend the RT range of the proteoform. Let *S*_1_, …, *S*_*i*-1_, *S*_*i*_, *S*_*i*+1_, …, *S*_*n*_ be all MS1 spectra in the increasing order of RT, in which the seed spectrum *S*_*i*_ contained the seed envelope *E*. We first checked if the spectrum *S*_*i-*1_ contained a matched experimental envelope of *E*. The isotopic peaks in *E* were matched to the experimental peaks in the spectrum to obtain an experimental envelope with an *m*/*z* error tolerance of 0.008. If two or more experimental peaks were matched to one theoretical peak, the one with the highest intensity was selected. Peaks in *E* were scaled to fit the peak intensities of the experimental peaks using the method described in Section 4.3. The scaled peaks in *E* with an intensity lower than the cutoff intensity of 3*h* (described in Section 4.2) were removed from the envelope along with the corresponding matched experimental peaks.

An experimental envelope was matched to the theoretical one if at least two of the three highest theoretical peaks matched experimental peaks. We searched for matched experimental envelopes in the neighboring spectra *S*_*i*-1_, …, *S*_1_ until we found two continuous spectra without a matched experimental envelope. The extension was also performed for the other direction in the neighboring spectra *S*_*i*+1_, …, *S*_*n*_.

The RTs of the first and last spectra reported by the extension method are called the initial start and end RTs of the proteoform, respectively. If a spectrum in the initial RT range contains a matched envelope, the corresponding trace intensity value is the sum of the intensities of the top three highest scaled theoretical peaks, and 0 otherwise. The trace intensities of all MS1 spectra in the initial RT range are called the extracted envelope chromatogram (XEC) of the seed envelope.

### 4.5 Adjusting RT boundaries

XECs of envelope sets were smoothed using a moving average filter with a window of size 2. Let *t*_*c*_ be the RT of a seed scan and *t*_s_ be the start RT of an envelope set. To adjust the start RT, we found all local minima in the XEC between *t*_*s*_ and *t*_*c*_ and ranked them in increasing order of intensity. Let *t*_*min*_ be the RT with the lowest XEC value *i*_*min*_. If there was a local maximum with RT *t*_max_ and trace intensity *i*_max_ such that *i*_max_ > 2.5*i*_min_ and *t*_min_ was between *t*_max_ and *t*_*c*_ (*t*_*s*_ < *t*_max_ < *t*_min_ < *t*_*c*_), then the start RT *t*_*s*_ was set to *t*_*min*_ (Supplementary Figure S6). The process was repeated until all the local minima had been checked. The process was performed to adjust the start and end RTs of reported envelope sets. This allowed us to fix errors in RT boundaries when the extended envelope set contained peaks from two or more neighboring envelope sets. All matched experimental envelopes in the adjusted RT range were reported as an envelope set of the proteoform.

### 4.6 Correcting charge states

Because of noise peaks, some seed envelopes reported from single spectra had an incorrect charge state. To correct change states, we summed up peak signals from several scans in an envelope set to obtain a better signal-to-noise ratio of peaks. For each peak in a seed envelope, the corresponding aggregate envelope peak was obtained by summing up the intensities of matched experimental peaks across all spectra within the RT range of the envelope set.

We used aggregated envelopes to fix one common type of error in charge states, in which a charge state *c* is mistakenly reported as charge state 2*c*. This type of error is called a double charge error. The main reason for double charge errors is that some noise peaks are randomly matched to theoretical peaks with charge state 2*c*.

The peaks in the aggregated envelope were ranked in the increasing order of their *m*/*z* values. The sums of even and odd index peaks were obtained for both theoretical and aggregate experimental envelopes. In an envelope with a double charge error, either all odd index or all even index peaks are from noise peaks, which tend to have low intensities. Let *A*_*e*_ and *A*_*o*_ be the sum of the intensities of even and odd aggregate experimental peaks, respectively. Let *B*_*e*_ and *B*_*o*_ be the sum of intensities of even and odd aggregate theoretical peaks, respectively. The two ratios *A*_*e*_*/B*_*e*_ *and A*_*o*_*/B*_*o*_ tend to be significantly different for envelopes with double charge errors. So, we calculated the log ratio with base 10 of the two ratios for each reported envelope set. If the absolute value of the log ratio was greater than 0.4, the charge state of the seed envelope was halved, and the new seed envelope was used to obtain an envelope set.

### 4.7 Extending envelope sets to envelope collections

After an envelope set with charge state *c* was reported, an envelope collection was obtained by exploring the neighboring charge states to find isotopic envelopes with the same monoisotopic mass. To find an envelope set with charge state *c*-1, the theoretical envelope *E*_*c-*1_ for charge state *c-*1 was obtained using the seed theoretical envelope of the envelope set with charge state *c*. Next, we extended *E*_*c-*1_ to obtain the start and end RTs using the methods in Section 4.4. If we failed to find at least two matched experimental peaks for the top three highest theoretical peaks in *E*_*c*-1_ in spectra *S*_*i*-1_, *S*_*i*_, and *S*_*i*+1_, then the envelope set for charge state *c*-1 was set to empty. We searched for envelope sets with charge states *c*-1, *c*-2, …,1 until two continuous empty envelope sets were found. Similarly, envelope sets were searched for charge states *c*+1, *c*+2 … until two continuous empty envelope sets were found. All identified non-empty envelope sets were added to the envelope collection.

### 4.8 Removing envelope collections from experimental data

To identify overlapping peaks shared by multiple envelope collections, we scaled peaks in a seed envelope to fit the peak intensities of its matched experimental envelope using the method in Section 4.3. If the intensity of an experimental peak was at least 4 times higher than that of the corresponding scaled theoretical peak, the peak was considered an overlapping one; otherwise, non-overlapping. To remove an envelope collection, the intensity of an overlapping experimental peak was reduced by the intensity of its matched theoretical peak, and non-overlapping experimental peaks were removed directly.

### 4.9 Refining monoisotopic masses of envelope collections

For an experimental peak *p* in an envelope collection, the *m*/*z* error between *p* and its matched theoretical peak is represented by *e*(*p*) and the intensity of its matched theoretical peak is represented by *h*(*p*). The weighted average *m*/*z* error of all peaks *p* in the envelope collection is 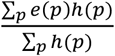, and the weighted average mass error of the envelope collection is the product of the average *m*/*z* error and the charge state of the seed envelope. The refined monoisotopic mass of an envelope collection was obtained by subtracting the weighted average error mass from its original monoisotopic mass.

### 4.10 Merging envelope collections

Once an envelope collection *F* was reported, we checked if it could be merged with another envelope collection. Two envelope collections were merged if (1) the difference between their masses was within [−1.00235 − *ϵ*, −1.00235 + *ϵ*], [−*ϵ, ϵ*], [1.00235 − *ϵ*, 1.00235 + *ϵ*], where *ϵ* was an error tolerance of 10 ppm and 1.00235 Da is an estimate of the mass difference of neighboring isotopic peaks in an envelope [38], (2) their RT ranges overlap was more than 80% of *F*, and (3) their change states ranges were not separated by more than 2 charge states.

### 4.11 Determining artifact masses

For a mass *x* and a maximum shifted mass of 10 neutrons, the set *X* of shifted and unshifted masses of *x* consists of 21 masses *x*+1.00235*d* for *d*=-10, -9, …,10, where 1.00235 Da is an estimated mass difference between two isotopologues introduced by a neutron [38]. A mass *y* is an isotopologue of mass *x* if *y* matches a mass in *X* with an error tolerance of 10 ppm. And *y* is a low (high) harmonic mass of *x* if the mass *yc* (*y*/*c*) matches a mass in *X* with an error tolerance of 10 ppm, where *c* is an integer.

### 4.12 The neural network model for ECScore

The neural network model for ECScore takes eight attributes of envelope collections as the input (Supplementary Table S7). The neural network model consists of four hidden layers (200 neurons in each layer) and an output layer. The activation function for the hidden layers is the Leaky Rectified Linear Unit with a negative slope coefficient of 0.05, and the sigmoid activation function for the output layer. L1 kernel regularization is applied to hidden layers with a regularization factor of 1×10^−6^. The neural network model was implemented using TensorFlow (version 2.7.0). In model training, the loss function was binary cross-entropy and the Adam optimizer with a learning rate of 1×10^−5^ was used. The training process was stopped if the validation loss did not improve for 30 epochs, and the model with the smallest validation loss was reported. To deal with the class imbalance problem in training data, class weighting by the inverse class frequency was used.

## 4. Code and data availability

TopFD is available as part of the TopPIC suite at https://github.com/toppic-suite/toppic-suite/releases/tag/v1.6_beta. The SW480 and SW620 data sets are available at the MassIVE repository (ID: MSV000090488). The data and Python scripts for training the ECScore model and evaluating the performance of the feature extraction tools are available at https://www.toppic.org/software/toppic/topfd_supplemental.html.

## Acknowledgments

This research was funded by NIH through the grants R01GM118470, R01GM125991, and R01CA247863.

## Author contributions

Z.Y., L.S. and X.L. designed the methods and experiments. L.S. generated the SW480 and SW620 data sets. A.R.B. and X.L. implemented and tested the methods and wrote the manuscript.

