## Supplementary Information for "TopFD - A Proteoform Feature Detection Tool for Top-Down Proteomics"

### Supplementary Tables

**Supplementary Table S1.** Summary of bottom-up feature detection tools

| | |
|---|---|
| msInspect [1] | msInspect identifies candidate peaks for feature detection by locating local maxima in each scan. Peaks that elute over several spectra in an LC-MS map are extracted. Using the peptide isotopic distribution, the co-eluting peaks are grouped to report a peptide feature. Kullback–Leibler divergence is used to evaluate the similarity between observed and experimental isotopic distributions. |
| centWave [2] | centWave uses each peak in experimental data as a candidate seed peak. If a peak in a spectrum is observed in neighboring spectra within a certain *m*/*z* range, these peaks are grouped to obtain a mass trace. A continuous wavelet transform is applied to each mass trace to determine its retention time boundaries. |
| MaxQuant [3] | In MaxQuant, candidate seed peaks are obtained by finding local intensity maxima in each spectrum. A Gaussian distribution is fitted to the seed peak and other peaks with similar *m*/*z* values in the scan to obtain an *m*/*z*-intensity curve. Afterward, these *m*/*z*-intensity curves are connected in neighboring scans based on their central positions to get the 3-dimensional retention time profile of the feature. Finally, deconvolution is performed based on 3-dimensional retention time profiles extracted from the LC-MS map. |
| Dinosaur [4] | Dinosaur collects centroided peaks with similar *m*/*z* values in consecutive spectra to build mass traces. Subsequently, deconvolution is performed to obtain peptide features. |
| DeepIso [5] | DeepIso is comprised of two deep-learning-based modules. The first module scans the experimental LC-MS map along the retention time axis and determines the charge state and retention time range of a feature. The second module scans the experimental LC-MS map along the *m*/*z* axis to group peaks that belong to the same isotopic distribution to report peptide features. |
| MSTracer [6] | MSTracer extracts mass traces and then groups mass traces whose local maxima are located at similar RT and whose intensities match the isotopic distribution of a peptide. A support vector regression model is employed to evaluate overlapping mass trace groups and report the best scoring one. A deep-learning model is used to report the quality score for each peptide feature. |

**Supplementary Table S2.** Envelope collections reported from the SW480 data set and the two breast cancer data sets

| Data | Number of Seed Envelopes | Number of Envelope Collections |
|---|---|---|
| SW480 colorectal cell line data – Replicate 1 | 471,330 | 20,218 |
| SW480 colorectal cell line data – Replicate 2 | 524,968 | 22,623 |
| SW480 colorectal cell line data – Replicate 3 | 520,505 | 20,731 |
| Basal-like breast cancer data – Replicate 1 | 67,730 | 1,785 |
| Basal-like breast cancer data – Replicate 2 | 65,058 | 1,664 |
| Basal-like breast cancer data – Replicate 3 | 68,751 | 1,820 |
| Basal-like breast cancer data – Replicate 4 | 69,213 | 1,863 |
| Basal-like breast cancer data – Replicate 5 | 63,547 | 1,684 |
| Basal-like breast cancer data – Replicate 6 | 65,159 | 1,823 |
| Luminal-B breast cancer data – Replicate 1 | 64,141 | 1,727 |
| Luminal-B breast cancer data – Replicate 2 | 64,466 | 1,902 |
| Luminal-B breast cancer data – Replicate 3 | 71,712 | 1,767 |
| Luminal-B breast cancer data – Replicate 4 | 67,956 | 1,833 |
| Luminal-B breast cancer data – Replicate 5 | 69,770 | 1,554 |
| Luminal-B breast cancer data – Replicate 6 | 68,126 | 1,758 |

**Supplementary Table S3.** Parameter settings for TopFD

| **Input Parameter** | **Value** |
|---|---|
| Maximum charge | 30 |
| Maximum mass | 70000 |
| MS1 signal noise ratio in MS-Deconv | 3.0 |
| *M*/*z* error tolerance in MS-Deconv | 0.02 |
| Disable final filtering in MS-Deconv | True |
| Use EnvCNN score in MS-Deconv | False |
| PCC cutoff for seed envelopes | 0.5 |
| Signal noise ratio for peak filtering | 3.0 |
| *M*/*z* error tolerance for peak filtering | 0.01 |
| *M/z* error tolerance for envelope extension | 0.008 |
| ECScore cutoff | 0.5* |

* ECScore cutoff was not used in the generation of training data sets for the neural network model

**Supplementary Table S4.** Parameter settings for ProMex

| **Parameter** | **Value** |
|---|---|
| Output mass range | Min 600, Max 100000 |
| Charge range | Min 1, Max 60 |
| Score threshold | -10 |

**Supplementary Table S5.** Parameter settings for FlashDeconv

| **Parameter** | **Value** |
|---|---|
| Minimum precursor signal noise ratio | 1 |
| mzML mass charge | 0 |
| Preceding MS1 count | 3 |
| Maximum MS level | 2 |
| Merging method | 0 |
| Tolerance | [10.0, 10.0] |
| Output mass range | Min 100, Max 100000 |
| Charge range | Min 1, Max 60 |
| *m*/*z* range | Min -1, Max -1 |
| RT range | Min -1, Max -1 |
| Minimum isotope cosine | [0.8, 0.8] |
| Minimum Q-score | 0 |
| Minimum peaks | [3, 3] |
| Minimum intensity | 100 |
| RT window | 180 |
| Mass error (ppm) | 10 |
| Mass error (Da) | -0.1 |
| Quant method | Area |
| Minimum sample rate | 0.2 |
| Minimum trace length | 1 |
| Maximum trace length | -1 |
| Minimum isotope cosine | -1 |

**Supplementary Table S6.** Parameter settings for Xtract

| **Parameter** | **Value** |
|---|---|
| Source spectrum type | Sliding windows |
| Use restricted time | Disabled |
| Chromatogram trace type | TIC |
| Sensitivity | High |
| Rel. Intensity threshold (%) | 1 |
| Target avg spectrum offset | 3 |
| Merge tolerance | 30 ppm |
| Max RT gap | 1 minute |
| Minimum number of detected intervals | 1 |
| Deconvolution algorithm | Xtract (isotopically resolved) |
| Output mass range | Min 100, Max 100000 |
| Output mass | M |
| Signal noise ratio threshold | 3 |
| Rel. Abundance threshold (%) | 0 |
| *m*/*z* range | Min 400, Max 2000 |
| Charge range | Min 1, Max 60 |
| Minimum number of detected charge states | 1 |
| Isotope table | Protein |
| Calculate XIC | Disabled |
| Fit factor (%) | 80 |
| Remainder threshold (%) | 25 |
| Consider overlaps | Enabled |
| Resolution at 400 m/z | RAW file specific |
| Negative charge | Disabled |
| Charge carrier | H+ (1.00727663) |
| Minimum intensity | 1 |
| Expected intensity error | 3 |

**Supplementary Table S7.** The total numbers of features, valid features, and mass artifacts reported from the OC and SW620 data sets by TopFD, ProMex, FlashDeconv, and Xtract

| Data | Tool | Number of Reported Features | Valid Features | Mass Artifacts |
|---|---|---|---|---|
| Ovarian Cancer – Replicate 1 | TopFD | 7459 | 7228 | 231 |
| | ProMex | 12018 | 5811 | 6207 |
| | FlashDeconv | 6346 | 6067 | 279 |
| | Xtract | 9319 | 7773 | 1546 |
| Ovarian Cancer – Replicate 2 | TopFD | 7755 | 7435 | 320 |
| | ProMex | 12043 | 5819 | 6224 |
| | FlashDeconv | 6133 | 5832 | 301 |
| | Xtract | 9672 | 7879 | 1793 |
| Ovarian Cancer – Replicate 3 | TopFD | 7865 | 7604 | 261 |
| | ProMex | 12640 | 6105 | 6535 |
| | FlashDeconv | 6336 | 6034 | 302 |
| | Xtract | 9969 | 8289 | 1680 |
| Ovarian Cancer – Replicate 4 | TopFD | 8144 | 7889 | 255 |
| | ProMex | 13184 | 6283 | 6901 |
| | FlashDeconv | 6632 | 6322 | 310 |
| | Xtract | 10324 | 8672 | 1652 |
| Ovarian Cancer – Replicate 5 | TopFD | 7957 | 7683 | 274 |
| | ProMex | 12694 | 6164 | 6530 |
| | FlashDeconv | 6442 | 6131 | 311 |
| | Xtract | 10061 | 8319 | 1742 |
| Ovarian Cancer – Replicate 6 | TopFD | 7804 | 7539 | 265 |
| | ProMex | 12633 | 6099 | 6534 |
| | FlashDeconv | 6312 | 6007 | 305 |
| | Xtract | 10057 | 8373 | 1684 |
| Ovarian Cancer – Replicate 7 | TopFD | 7799 | 7551 | 248 |
| | ProMex | 12734 | 6109 | 6625 |
| | FlashDeconv | 6337 | 6055 | 282 |
| | Xtract | 10388 | 8609 | 1779 |
| Ovarian Cancer – Replicate 8 | TopFD | 8002 | 7765 | 237 |
| | ProMex | 13166 | 6440 | 6726 |
| | FlashDeconv | 6464 | 6129 | 335 |
| | Xtract | 10562 | 8643 | 1919 |
| Ovarian Cancer – Replicate 9 | TopFD | 8069 | 7782 | 287 |
| | ProMex | 13213 | 6368 | 6845 |
| | FlashDeconv | 6379 | 6074 | 305 |
| | Xtract | 10660 | 8849 | 1811 |
| Ovarian Cancer – Replicate 10 | TopFD | 7961 | 7693 | 268 |
| | ProMex | 13125 | 6366 | 6759 |
| | FlashDeconv | 6300 | 6004 | 296 |
| | Xtract | 10852 | 8931 | 1921 |
| SW620 colorectal cell line data – Replicate 1 | TopFD | 12764 | 10576 | 2188 |
| | ProMex | 6264 | 3025 | 3239 |
| | FlashDeconv | 15801 | 12240 | 3561 |
| | Xtract | 10927 | 8322 | 2605 |

| | | | | |
|---|---|---|---|---|
| SW620 colorectal cell line data – Replicate 1 | TopFD | 12578 | 10410 | 2168 |
| | ProMex | 6090 | 2984 | 3106 |
| | FlashDeconv | 16521 | 12781 | 3740 |
| | Xtract | 11046 | 8379 | 2667 |
| SW620 colorectal cell line data – Replicate 1 | TopFD | 13598 | 11145 | 2453 |
| | ProMex | 6320 | 3136 | 3184 |
| | FlashDeconv | 16663 | 13034 | 3629 |
| | Xtract | 11968 | 9101 | 2867 |

**Supplementary Table S8.** Eight input attributions of envelope collections in the neural network model for ECScore

| Attribute | Description |
|---|---|
| EnvCNN Score | EnvCNN score of the aggregate envelope obtained from the envelope collection (see Section 2.3) |
| Scaled retention time range | Retention time range (in minutes) of the envelope set of the seed envelope scaled by 1/60 |
| Ratio of matched peaks | Ratio between the total number of matched experimental peaks and the total number of theoretical peaks |
| Total peak intensity with log transformation | Logarithm of the sum of the scaled intensities of all theoretical peaks |
| Seed charge state | Charge state of the seed envelope |
| Average correlation of three experimental envelopes | Three experimental envelopes with the seed charge state are extracted from the spectrum of the seed and two neighboring spectra. Pearson's correlation is computed for each pair of the three envelopes and the average correlation of the three pairs is reported. |
| Scaled charge state range | Charge state range (maximum charge – minimum charge +1) scaled by 1/30 |
| Log ratio for charge state correction | The log ratio for charge state correction is computed using the method in Section 4.6. |

### Supplementary Figures

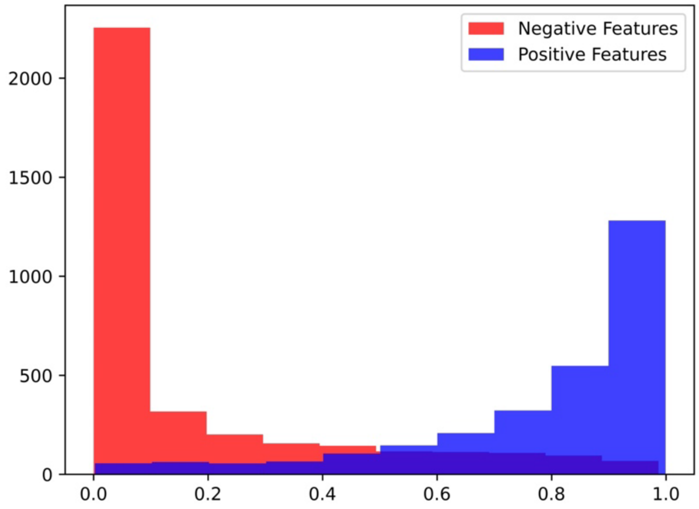

**Supplementary Figure S1.** Histograms of ECScores of positive and negative proteoform features (envelope collections) in the validation data set generated from the SW480 and two breast cancer data sets.

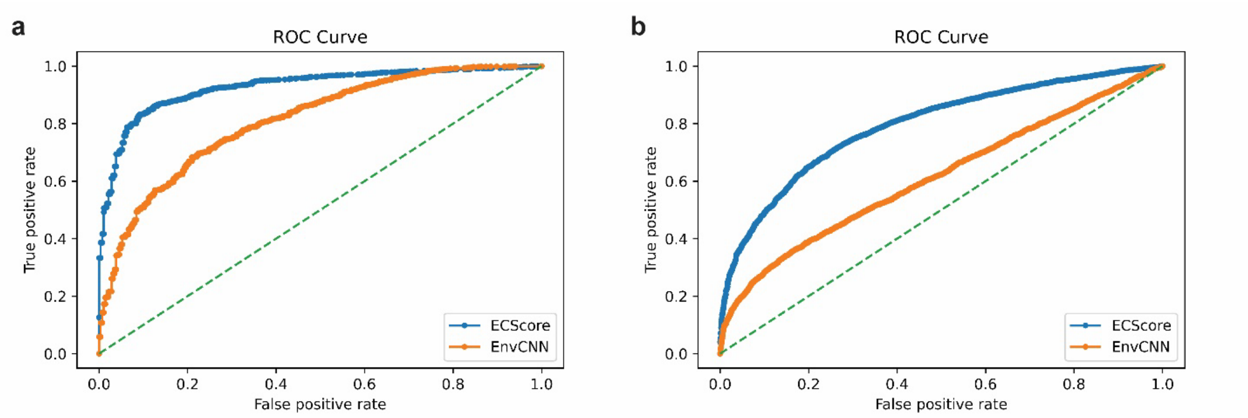

**Supplementary Figure S2.** Comparison between ECScore and the EnvCNN score on the OC and SW620 test data. (a) ROC curves on the OC test envelope collections. (b) ROC curves on the SW620 test envelope collections.

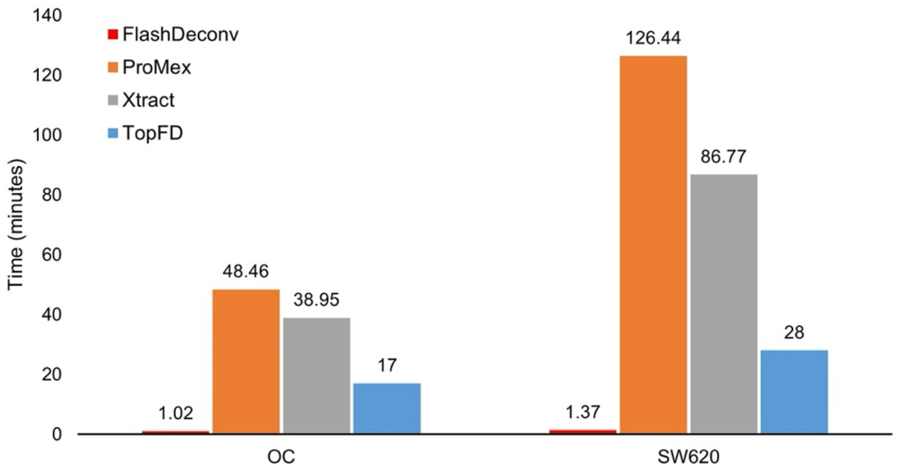

**Supplementary Figure S3.** Running times of TopFD, ProMex, Xtract, and FlashDeconv on the first OC replicate and the first SW620 replicate. The running time of each tool was obtained on a desktop computer with an Intel® Core™ i7-8700 @ 3.2GHz CPU and 16 GB RAM using 1 CPU thread. Only MS1 spectra were deconvoluted and MS/MS spectra were not deconvoluted in the test.

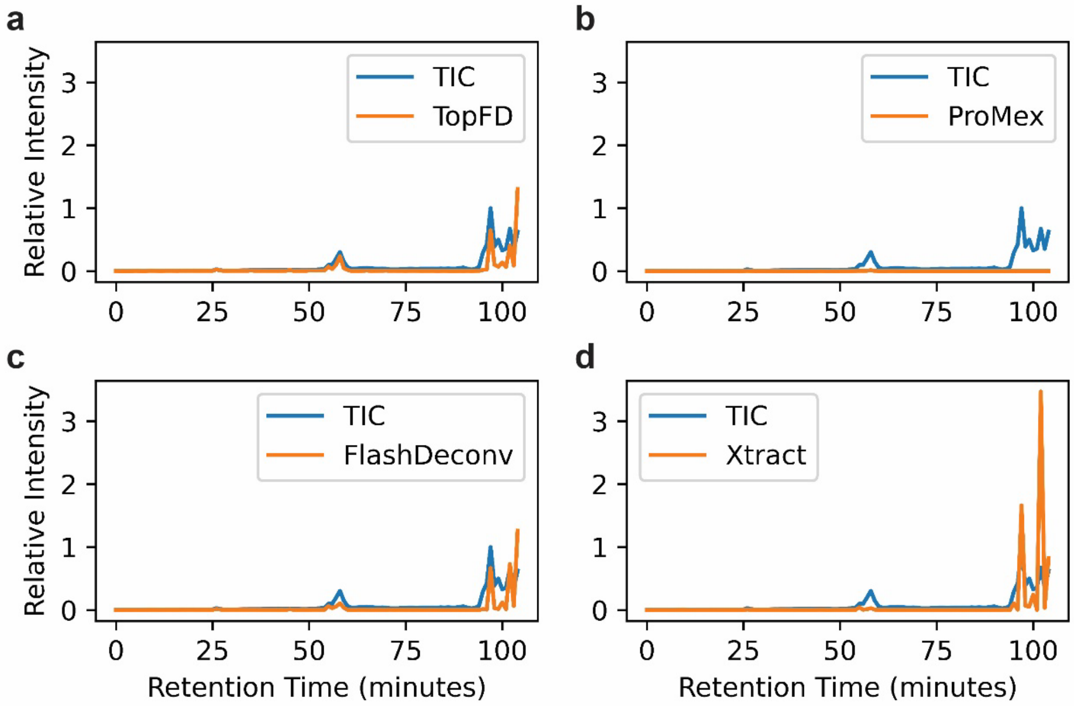

**Supplementary Figure S4.** Comparison of TICs and total proteoform feature intensities reported by four feature detection tools along the RT time for the first SW620 replicate. (a) TopFD, (b) ProMex, (c) FlashDeconv, and (d) Xtract. The RT range of the MS data is divided into 1-minute RT bins. The TICs and total proteoform feature intensities are normalized by dividing them by the maximum TIC value.

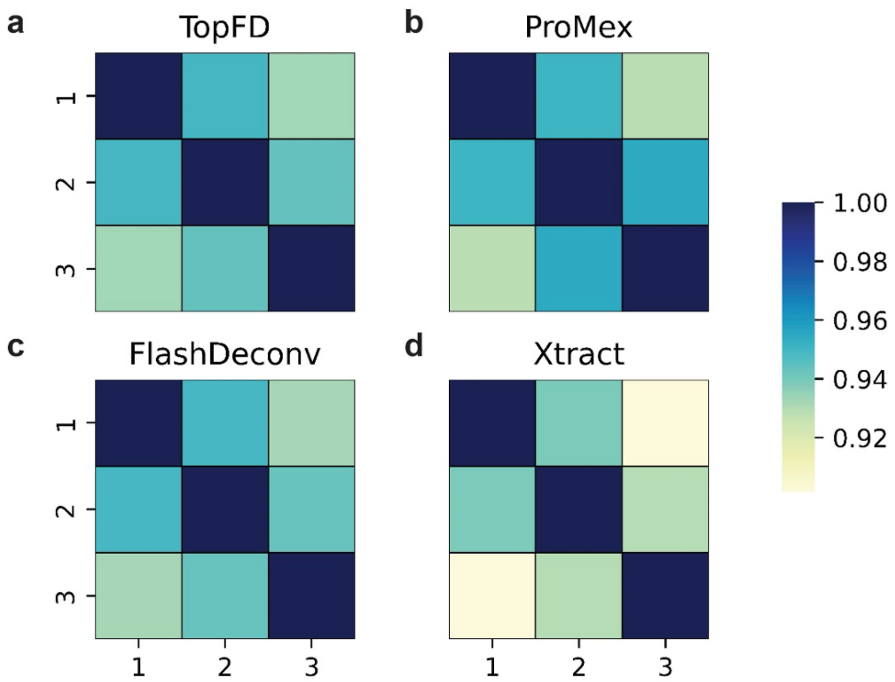

**Supplementary Figure S5.** Quantitative reproducibility of proteoform features reported from the SW620 data set. The PCC between the log-abundances of proteoform features is obtained for each replicate pair for a) TopFD, b) ProMex, c) FlashDeconv, and d) Xtract.

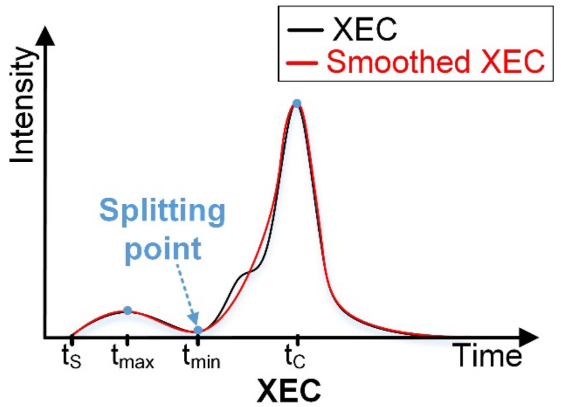

**Supplementary Figure S6.** An illustration of adjusting the RT boundaries of an envelope set. The XEC of the envelope set contains peaks from two neighboring envelope sets. The XEC is smoothed by employing a moving average filter with a window of 2. $t_C$ is the RT of the spectrum with the seed envelope, and $t_S$ is the original start RT of the envelope set. A local minimum is located at $t_{min}$ and a local maximum is located at $t_{max}$. The intensity ratio of XEC at $t_{max}$ and $t_{min}$ is greater than 2.5, so the start RT is adjusted to $t_{min}$.
